# Deficits in integrative NMDA receptors caused by *Grin1* disruption can be rescued in adulthood

**DOI:** 10.1101/2022.11.22.517556

**Authors:** S. Venkatesan, M.A. Binko, C.A. Mielnik, A.J. Ramsey, E.K. Lambe

## Abstract

Glutamatergic NMDA receptors (NMDAR) are critical for cognitive function, and their reduced expression leads to intellectual disability. Since subpopulations of NMDARs exist in distinct subcellular environments, their functioning may be unevenly vulnerable to genetic disruption. Here, we investigate synaptic and extrasynaptic NMDARs on the major output neurons of the prefrontal cortex in mice deficient for the obligate NMDAR subunit encoded by *Grin1* and wild-type littermates. With whole-cell recording in brain slices, we find that single, low-intensity stimuli elicit surprisingly-similar glutamatergic synaptic currents in both genotypes. By contrast, clear genotype differences emerge with manipulations that recruit extrasynaptic NMDARs, including stronger, repetitive, or pharmacological stimulation. These results reveal a disproportionate functional deficit of extrasynaptic NMDARs compared to their synaptic counterparts. To probe the repercussions of this deficit, we examine an NMDAR-dependent phenomenon considered a building block of cognitive integration, basal dendrite plateau potentials. Since we find this phenomenon is readily evoked in wild-type but not in *Grin1*-deficient mice, we ask whether plateau potentials can be restored by an adult intervention to increase *Grin1* expression. This genetic manipulation, previously shown to restore cognitive performance in adulthood, successfully rescues electrically-evoked basal dendrite plateau potentials after a lifetime of NMDAR compromise. Taken together, our work demonstrates NMDAR subpopulations are not uniformly vulnerable to the genetic disruption of their obligate subunit. Furthermore, the window for functional rescue of the more-sensitive integrative NMDARs remains open into adulthood.

## Introduction

Glutamatergic N-methyl-D-aspartate receptors (NMDARs) are increasingly appreciated for their role in cognitive integration ^1-4^. Mutations that reduce expression or function of NMDARs are a direct cause of intellectual disability ^5,6^. Relatively little is known, however, about whether there is variability across cellular domains in the functional impact of NMDAR genetic compromise. This is a critical area of exploration because NMDARs in different subcellular compartments play distinct neurophysiological roles ^2,7,8^ and experience distinct regulatory environments that may permit differing degrees of homeostatic compensation ^9-14^. Understanding the relative vulnerability of NMDAR subpopulations to genetic disruption is essential to appreciate mechanisms of cognitive compromise and to identify new treatment approaches.

NMDARs are high affinity ligand-gated channels that are also voltage-dependent, requiring both ligand-binding and depolarization to open. If glutamate binds without sufficient depolarization to relieve Mg^2+^ blockade, the NMDAR acts as a ‘coincidence-detector’ between synaptic activation and subsequent depolarization. While this concept has been well explored in the context of synaptic plasticity, it is increasingly appreciated that glutamate travels beyond the synapse and this spillover increases upon strong or repeated stimuli ^15-17^. Glutamate spillover is the substrate for integrative phenomena including dendritic plateau potentials, where stimulation of extrasynaptic NMDARs in the healthy brain allows enhanced cortical output in response to strong, repetitive, or converging inputs ^2,7,18-20^.

Here, we investigated *Grin1* knockdown (*Grin1*KD) mice with a profound deficiency in NMDAR receptor expression and binding ^21,22^ and severe cognitive deficits ^23^. Consistent with previous work in this mouse and in other models of developmental cognitive disruption ^24^, neuronal membrane properties are unaltered. Furthermore, low-intensity stimuli revealed that neither AMPA receptor (AMPAR) nor NMDAR synaptic currents differed significantly across genotypes. However, a sizable deficit in the *Grin1*KD NMDAR response was revealed by stronger, repetitive, or pharmacological stimulation. The magnitude of this functional deficit was consistent with deficits observed anatomically in previous receptor binding work. To probe the repercussions of this primarily extrasynaptic deficit in NMDARs, we examined dendritic plateau potentials and found that *Grin1*KD mice are severely impaired in this integrative domain. In the final experiment, we tested the possibility of restoring cognitively-critical synaptic integration in adulthood, building on recent work showing that adult intervention to increase *Grin1* expression achieves meaningful cognitive restoration ^23^. We determine that dendritic plateau potentials can indeed be rescued by adult intervention to increase *Grin1* expression. Taken together, this work reveals that integrative NMDARs are disproportionately sensitive to genetic disruption but amenable to restoration upon intervention in adulthood.

## Materials and Methods

### Animals

All experiments were approved by the University of Toronto Animal Care and Use Committee and followed Canadian Council on Animal Care guidelines. Mice were group-housed and kept on a 12-hour light cycle with food and water access *ad libitum*.

Mice for the initial experiments were generated from intercross breeding of C57Bl/6J *Grin1* heterozygotes with 129X1Sv/J *Grin1* heterozygotes, producing an F1 generation of *Grin1*KD (*Grin1*^neo/neo^) and wild-type (WT) littermate siblings used for experiments ^21,23^. Adult male and female mice were used for experiments (sex-matched and age-matched; age: 104 ± 5 days), with recordings from 52 WT and *Grin1*KD mice.

For the adult genetic rescue experiments, we used an additional 14 WT, *Grin1KD*, and *Grin1*rescue mice of both sexes. The generation of the line permitting adult rescue with tamoxifen is described in greater detail ^23^. Starting in adulthood at 84 ± 6 days, all three genotypes of mice for the rescue experiment were treated with tamoxifen chow (TD.140425, 500□mg/kg, Envigo) *ad libitum* for 14 days. Electrophysiology experiments were conducted upon 38 ± 5 days washout from tamoxifen (sex matched and age-matched; age:135 ± 3 days).

### Electrophysiological Recordings

Prefrontal brain slices were prepared as previously described ^25^ and as detailed in **Supplemental Methods**. Layer 5 pyramidal neurons in the medial prefrontal cortex, including cingulate and prelimbic regions, were visually identified by their pyramidal shape and prominent apical dendrite using infrared differential inference contrast microscopy. Unless otherwise indicated, whole-cell patch clamp electrodes contained potassium-gluconate patch solution. All ACSF and pipette solutions used for the following experiments are listed in **Supplemental Methods**. Intrinsic membrane properties and excitability were assessed in current-clamp.

### Evoked excitatory postsynaptic currents

AMPAR-mediated evoked excitatory postsynaptic currents (eEPSCs) were measured in voltage-clamp at a holding potential of −75 mV. A bipolar stimulating electrode (FHC) was located in layer 2/3 for apical dendrite stimulation with pyramidal neurons in layer 5 recorded ∼250 μm away from the electrode. For basal dendrite stimulation, the stimulating electrode was placed in the basal dendritic field ∼100 μm from the soma of the recorded layer 5 pyramidal neuron. For both apical and basal stimulation paradigms, single pulses of 40 μs duration were delivered at 0.1 Hz, increasing in 10 μA increments. The AMPAR–mediated eEPSCs were analyzed as an average of at least 3 traces with Clampfit (Molecular Devices).

Isolated NMDA receptor–mediated eEPSCs were measured in voltage-clamp at a holding potential of +60 mV using specialized patch solution to block voltage-gated potassium and sodium channels. These recordings were performed in the presence of modified ACSF (1 mM MgSO_4_), AMPAR antagonists CNQX (20 μM) or NBQX (20 μM), and GABA receptor antagonists picrotoxin (PTX, 50 μM) and CGP52432 (CGP, 1 μM). Stimulation in the apical or basal dendritic fields were delivered as above. The NMDA receptor–mediated eEPSCs were analyzed as an average of 3 traces with Clampfit (Molecular Devices) and D-APV (50 μM) was applied to confirm NMDAR responses.

### Enhancing glutamate spillover

To additionally recruit the extrasynaptic population of NMDA receptors, a 20 Hz train of mild stimuli was delivered in the apical location. Glutamate spillover was additionally enhanced with the application of TBOA (30 μM) and LY341495 (1 μM) to block glial glutamate uptake and mGluR2/3 presynaptic autoreceptors respectively ^26,27^.

### Pharmacological stimulation with NMDA application

Total synaptic and extrasynaptic NMDAR currents were measured by bath application of NMDA (20 μM, 30 s) in a different subset of brain slices. Voltage-clamp recordings were performed with potassium-gluconate patch solution in a modified ACSF to reduce magnesium blockade as neurons were held at –75 mV. The AMPAR antagonist CNQX (20 μM) was also included. The peak amplitude of the NMDA receptor current was compared to baseline current using Clampfit. In a subset of experiments, D-APV (50 μM) was applied to verify NMDAR mediation of the inward currents.

### NMDAR-dependent dendritic plateau potentials

Plateau potentials were generated by stimulation of the basal dendritic field of layer 5 pyramidal neurons, with the stimulating electrode placed within ∼100 μm radius of the cell body. Plateau potentials were recorded in current-clamp at an initial membrane potential of −75 mV. They were generated with 10 stimuli at 50 Hz at the minimal stimulus intensity to evoke glutamatergic EPSCs ^7,28^. PTX (20 μM) and CGP52432 (1 μM) were present to block GABA receptors in combination with AMPAR blockers CNQX (20 μM) or NBQX (20 μM) to isolate NMDAR plateau potentials. D-APV (50μM) was applied to confirm NMDAR dependence of plateau potentials.

### Statistics

Statistical tests were performed in Prism 7 (Graphpad). Data are presented as mean ± SEM. Parametric statistical comparisons between responses from different groups of mice were determined using two-tailed unpaired *t* tests, and within-cell effects examined with two-tailed paired *t* tests. Where appropriate, interactions between genotype and other variables were assessed with two-way ANOVA or repeated-measure two-way ANOVA with *post hoc* Sidak-corrected *t* tests. Where 3 groups were treated with tamoxifen, the impact of adult intervention to rescue *Grin1* expression was assessed with non-parametric Kruskal Wallis test and Dunn’s post hoc tests due to the distribution of the data. Within cell pharmacological investigations for this dataset were therefore compared with a non-parametric paired test.

## Results

To investigate the differential vulnerability of synaptic and extrasynaptic NMDARs to genetic disruption, we performed *ex vivo* electrophysiology in major output pyramidal neurons of prefrontal cortex from mice deficient in the obligate NMDAR subunit (*Grin1*KD) and their wild-type (WT) littermate controls (**Fig 1A**). We found that neuronal properties, including resting membrane potential, input resistance, capacitance, spike amplitude, and rheobase did not differ significantly between the genotypes (**Supplemental Table S1**). The input-output relationship showed the expected effect of current (*F*_3,123_ = 307.6; *P*<0.0001; **Fig 1B,C**), but did not differ significantly between the genotypes (*F*_1, 41_ = 0.4525; *P*=0.50), nor show an interaction *F*_3,123_ = 1.123; *P*=0.34).

**Figure 1.**
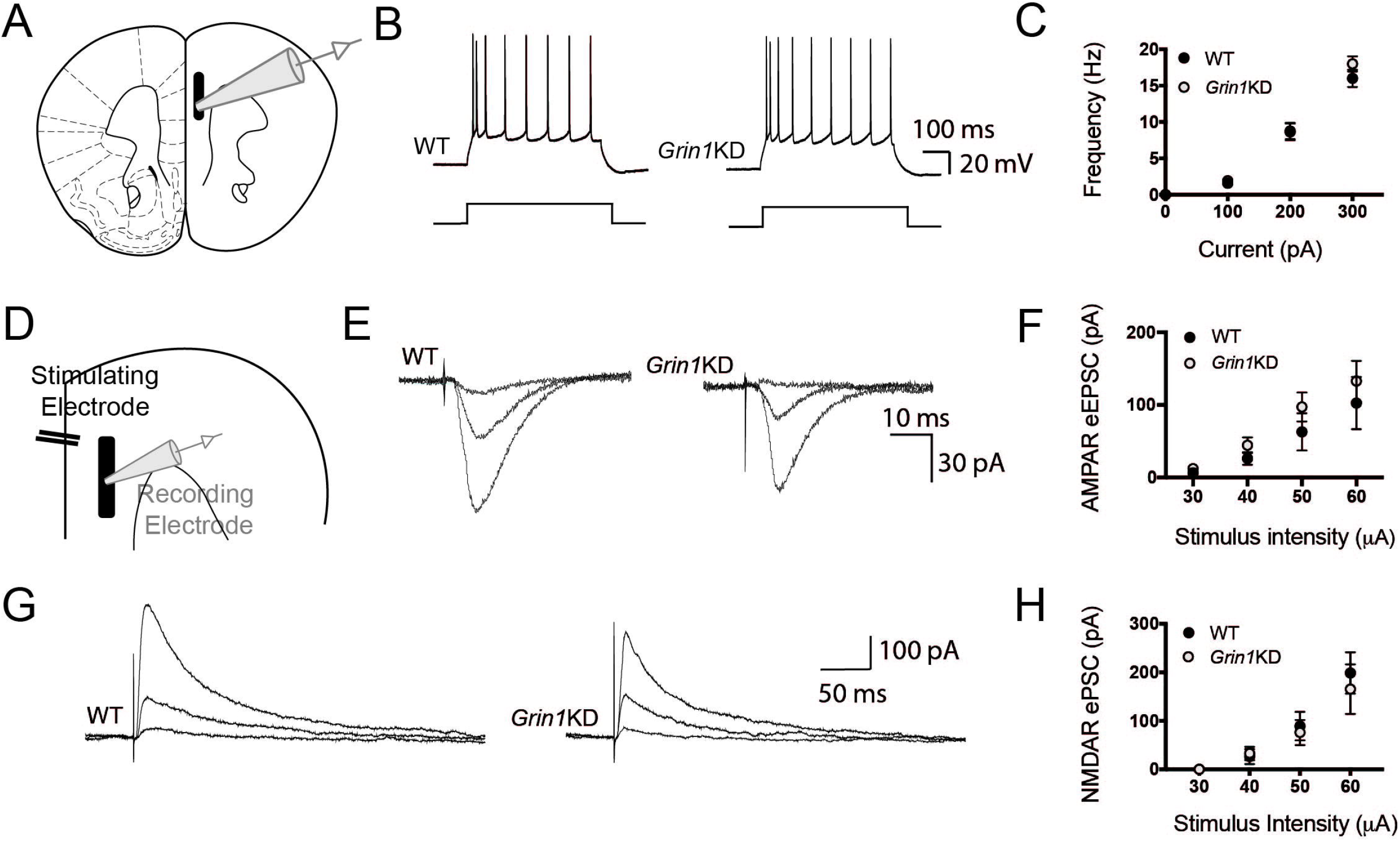
Wild-type and *Grin1*KD have similar intrinsic excitability and postsynaptic AMPA and NMDA receptor responses. (**A**) Schematic of the prefrontal cortex with electrophysiological recording from layer 5 pyramidal neuron. (**B**) Example current-clamp traces from WT (left) and *Grin1*KD (right) in response to depolarizing current steps through the recording pipette. (**C**) Input-output graphs of spike frequency (Hz) in current-clamp for WT (*n* = 20) and *Grin1*KD (*n* = 25). (**D**) Schematic of recording pipette with extracellular stimulating electrode for assessment of postsynaptic currents. (**E**) Example voltage-clamp traces (Vh −75 mV) show inward AMPA receptor (AMPAR)-mediated electrically-evoked excitatory postsynaptic currents (eEPSC) in WT and *Grin1*KD. (**F**) Graph illustrates that WT (*n* = 15) and *Grin1*KD (*n* = 13) both show the expected relationship between stimulus strength eEPSC amplitude but no significant effect of genotype nor interaction for AMPAR eEPSCs. (**G**) Example voltage-clamp traces (Vh +60 mV) show outward NMDA receptor (NMDAR)-mediated evoked postsynaptic currents (ePSCs), isolated with AMPAR and GABA receptor blockade and recorded with pipette solution to internally block voltage-gated potassium and sodium channels. (**H**) Graph illustrates that both WT (*n* = 10) and *Grin1*KD (*n* = 15) show the expected relationship between stimulus strength and NMDAR ePSC amplitude, but no significant effect of genotype nor interaction for these ePSCs. Data is represented as mean ± SEM.

### Preserved synaptic glutamatergic responses in Grin1KD mice

To test AMPAR synaptic responses from stimulation in the apical dendritric field, we recorded from layer 5 pyramidal neurons at a holding potential of −75 mV and applied electrically-evoked stimulation in layer 2/3 (**Fig 1D**). There was no significant difference between genotypes in the electrical stimulus required to elicit the minimal response (*t*_27_= 0.3; *P* = 0.8), and response amplitudes were similar in both genotypes (**Fig 1E,F)**. We observed the expected effect of stimulus strength on response amplitude (*F*_3,115_ = 11.04; *P*<0.0001), but not an effect of genotype (*F*_1,115_ = 2.354; *P* = 0.13), nor an interaction between genotype and stimulus strength (*F*_3, 115_ = 0.20; *P* = 0.9). These results show that AMPAR-mediated synaptic transmission in response to low-intensity stimulation is similar in WT and *Grin1*KD prefrontal cortex.

To isolate NMDAR synaptic responses, we next recorded evoked currents at a holding potential of +60 mV in the presence of AMPAR and GABA_A_ receptor antagonists, using recording pipette solution designed to block voltage-gated potassium and sodium channels. Again, there was no genotype difference in the minimal current required to elicit a response (*t*_24_ = 0.4; *P* = 0.71), nor in response amplitudes across an increasing range of stimuli (**Fig 1G,H)**. We observed the expected effect stimulus strength on response amplitude (*F*_3, 87_ = 12.53; *P*<0.0001), but no effect of genotype (F_1, 87_ = 0.1926; P=0.66), nor an interaction between genotype and stimulus strength (F_3, 87_ = 0.1485; P=0.93). Consistent with the intended NMDAR-mediation of these EPSCs, the evoked currents were strongly suppressed by the selective NMDAR antagonist, D-APV (50 μM): *t*_(10)_ = 6.1, *P* = 0.0001). These results demonstrate that the amplitudes of isolated NMDAR currents are similar between genotypes in response to low-intensity stimulation. This unexpected finding was surprising because of the prominent differences in the expression of the obligate subunit and NMDAR binding between the genotypes in previous reports ^21,22^.

We therefore hypothesized that deficits are more prominent in the extrasynaptic NMDAR subpopulation, which can be recruited by stronger electrical stimulation to increase glutamate spillover ^29,30^. Therefore, we delivered stronger single stimuli (80 μA) in a subsequent experiment. In contrast to the relatively-homogenous effects of low-intensity stimulation, stronger stimuli elicited a significant and substantial difference in NMDAR ePSC amplitude between genotypes (WT: 599 ± 105 pA, *n* = 9; Grin1KD: 339 ± 62 pA, *n* = 16; *t*_23_= 2.29; *P* = 0.032; data not shown). This result prompted a detailed characterization of extrasynaptic NMDAR in *Grin1*KD mice using multiple approaches.

### Deficient extrasynaptic NMDAR responses in Grin1KD mice

To recruit extrasynaptic NMDARs by boosting glutamate spillover, repetitive stimuli in a 20 Hz train were delivered under baseline conditions and then under standard conditions to increase glutamate spillover ^26,27^, suppression of glutamate reuptake with TBOA and autoinhibition with LY341495 (**Fig 2A,B**). In wild-type mice, repetitive stimulation led to summation of postsynaptic responses, yielding a higher peak response compared to the first input, with further potentiation of peak response caused by glutamate spillover in the presence of TBOA. In *Grin1*KD, by contrast, boosting spillover did not increase the peak response, leading to a significant interaction between the genotype and spillover conditions (*F*_2,16_ = 11.37; *P* = 0.0008). Repetitive stimulation in the presence of TBOA significantly potentiated the peak response compared to the first stimulus in WT (Sidak’s post hoc test, *P* = 0.0001) but not in *Grin1*KD (*P* = 0.2). These results suggest a lack of extrasynaptic NMDARs in *Grin1*KD available to be recruited by glutamate spillover.

**Figure 2.**
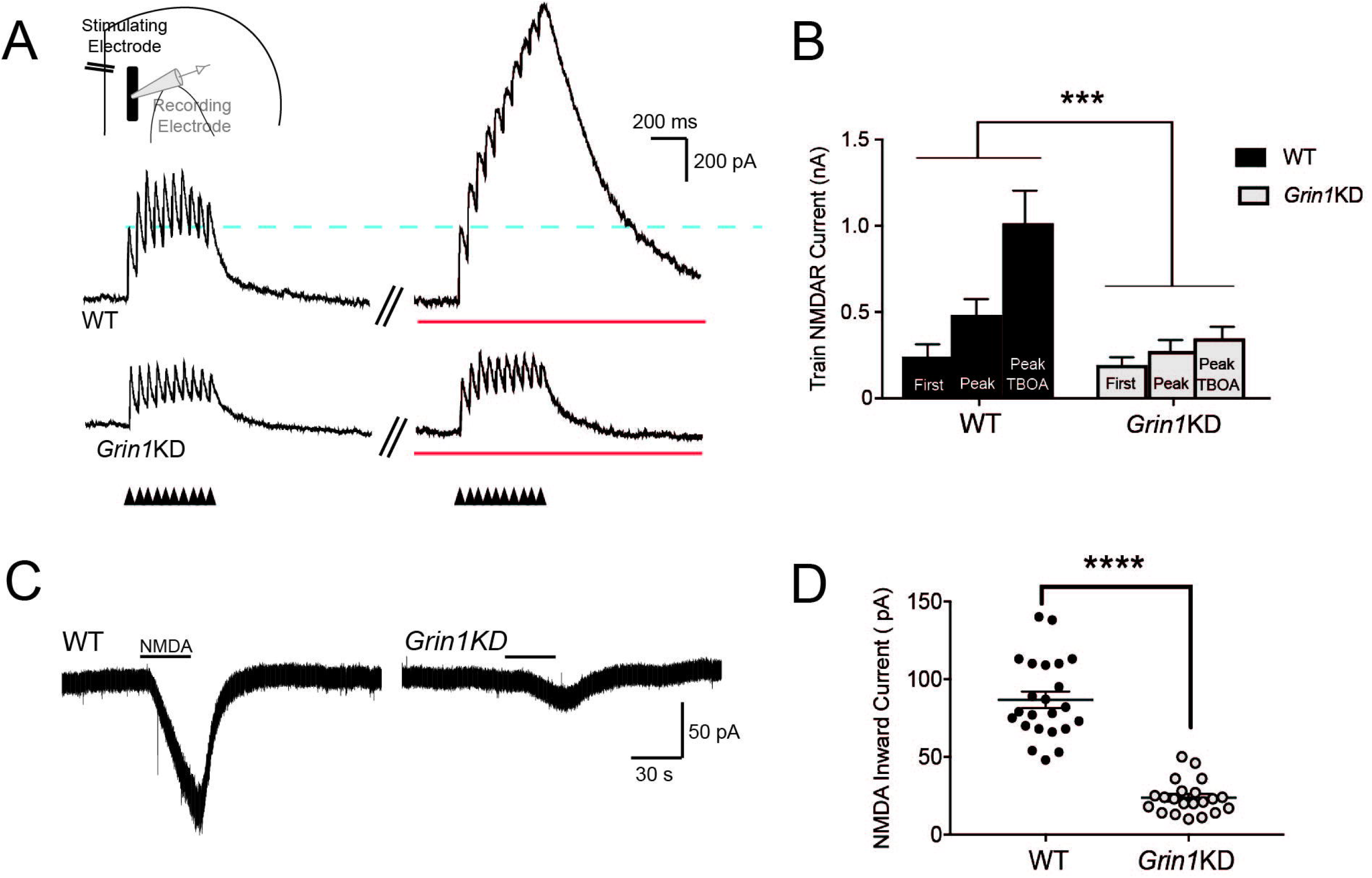
Extrasynaptic NMDARs are not recruited in *Grin1*KD mice during glutamate spillover. (**A**) Voltage-clamp traces (Vh +60 mV) show NMDAR-mediated outward currents during AMPAR blockade in WT (above) and *Grin1*KD (below) evoked by a stimulus train (20 Hz, 10 pulses) under baseline conditions and with the addition of TBOA and LY341495 to enhance glutamate spillover (red line). The dotted line illustrates the consistency of the first evoked postsynaptic current. NMDAR responses isolated with AMPAR and GABA receptor blockade. (**B**) The bar graph shows the significant potentiation of the peak amplitude in the stimulus train under conditions of enhanced glutamate spillover for WT (*n* = 4) but not *Grin1*KD (*n* = 6); significant interaction of genotype and spillover condition (\*\*\**P* < 0.001). (**C**) Voltage-clamp traces show bath application of NMDA to pharmacologically stimulate NMDAR in WT (left) and *Grin1*KD (right). (**D**) The bar graph shows the peak amplitude of pharmacologically-elicited inward NMDA currents is significantly lower in *Grin1*KD (*n* = 21) compared to WT (*n* = 23) (\*\*\*\**P* ≤ 0.0001). Data represented as mean ± SEM.

To reach an even broader group of extrasynaptic receptors, we activated NMDARs using direct pharmacological manipulation with the agonist NMDA. For these experiments, we bath-applied NMDA to the prefrontal slice in the presence of AMPAR antagonist CNQX and low-Mg^2+^ to permit NMDAR activation at a holding potential of −75 mV. As anticipated ^23^, pharmacological NMDAR currents were substantially and significantly reduced in *Grin1*KD mice compared to their littermates (WT: 87 ± 5 pA, *n* = 23; *Grin1*KD1: 24 ± 2 pA, *n* = 21; *t*_42_ = 10.6, *P* = 0.0001; **Fig 2C,D**). These pharmacologically-elicited inward currents were suppressed by D-APV (50 μM; WT: *n* = 5, *t*_4_ = 6.2, *P* = 0.003; *Grin1*KD mice: *n* = 7, *t*_6_ = 3.5, *P* = 0.01). Of note, the 3-fold genotype difference in the response to bath NMDA mirrors the difference in NMDAR binding observed in prefrontal cortex in *Grin1*KD compared to wild-type controls ^23^.

Stronger, repetitive, and pharmacological stimulations that recruit extrasynaptic NMDARs all unmask genotype differences between the wild-type littermates and *Grin1*KD mice, consistent with the interpretation that *Grin1*KD mice have a specific and disproportionate deficit in extrasynaptic NMDARs.

### Impact of extrasynaptic NMDAR disruption: Dendritic plateau potentials

Dendritic plateau potentials can be evoked by spillover of glutamate onto extrasynaptic NMDARs under conditions of high-frequency repetitive stimulation of inputs to basal dendrites ^7,28^. This integrative phenomenon depends on the recruitment of extrasynaptic NMDARs (**Fig 3A**), and would be vulnerable if this population were compromised (**Fig 3B**). Dendritic plateau potentials are considered an important cognitive substrate to link multiple streams of incoming information and generate burst firing ^16,19,20,31^, an output signal predicted to exert stronger downstream consequences ^32,33^. Deficient extrasynaptic NMDARs are predicted to have profound consequences for such signaling ^7,28^.

**Figure 3.**
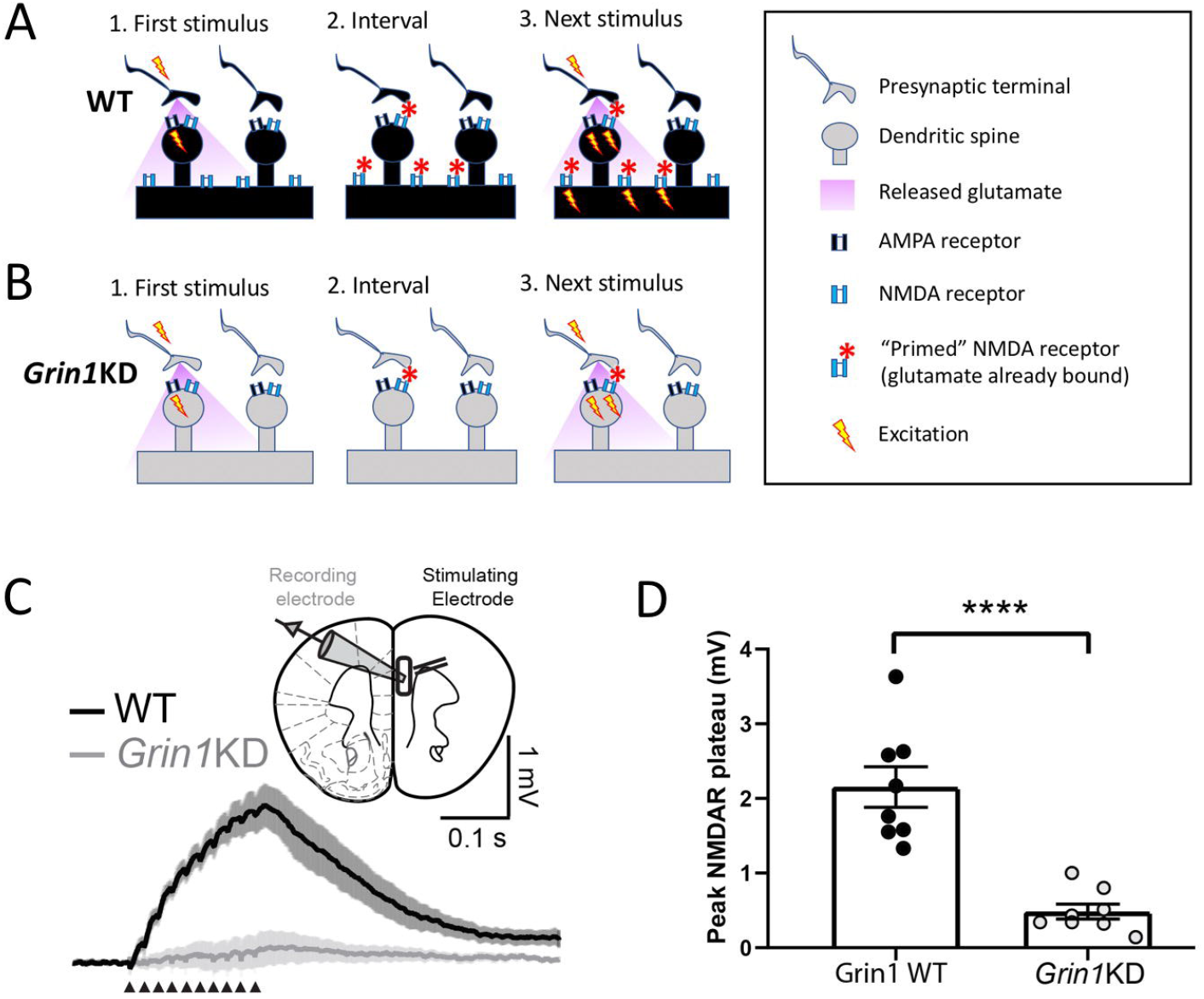
Deficits in extrasynaptic NMDA receptors disrupt integrative basal dendrite plateau potentials in *Grin1*KD. Schematics depict hypothesized differences in extrasynaptic NMDA receptors (NMDARs) between (**A**) WT and (**B**) *Grin1*KD. The initial stimulus (***1***.) yields glutamate spillover that permits priming of extrasynaptic NMDARs during the inter-stimulus interval (***2***.) making them available to be activated immediately by depolarization from the next stimulus (***3***.). This form of integration is sufficient to yield a dendritic plateau potential in response to repeated mild stimulation and is typically measured in current-clamp. (**C**) Inset: Schematic of layer 5 pyramidal cell recording with stimulation in the basal dendritic field. Averaged current-clamp recordings of excitatory responses to repeated minimal stimulation (50 Hz, 10 pulses, 30-40 μA) in WT (black, *n* = 8) and *Grin1*KD (gray, *n* = 8). NMDAR-mediated dendritic plateaus isolated with AMPA and GABA receptor blockade. (**D**) Graph of peak plateau amplitude illustrates that basal dendrite integration is substantially reduced in *Grin1*KD mice compared to WT (**** *P* < 0.0001). Data represented as mean ± SEM.

To examine basal dendrite plateau potentials in both genotypes, we recorded from layer 5 pyramidal neurons while electrically stimulating inputs in the basal field. AMPAR eEPSCs evoked by basal dendritic stimulation were similar between wild-type and *Grin1*KD mice, had the expected effect of current (F_2,66_ = 12.7; *P* = 0.0001), but no effect of genotype (*F*_1, 66_ = 0.148, *P* = 0.7) nor interaction between genotype and current (*F*_2, 66_ = 0.127, *P* = 0.88, data not shown). Next, we recorded NMDAR plateau potentials in current-clamp in response to trains of stimuli (50 Hz, 10 pulses) in the presence of AMPA and GABA receptor blockade and observed a marked genotype difference (**Fig 3C,D**). While wild-type neurons showed clear NMDAR plateau potentials (peak amplitude: 2.15 ± 0.27 mV, *n* = 8), the train of stimuli did not elicit dendritic plateau potentials in *Grin1*KD neurons (0.48 ± 0.10 mV, *n* = 8; *t*_14_ = 5.8 *P* < 0.0001; **Fig 3C,D**). Plateau potentials in wild-type neurons could be eliminated by the NMDAR antagonist APV (significant genotype x D-APV interaction: *F*_1, 7_ = 7.53, *P* = 0.029; peak amplitude at baseline vs APV in WT: *t*_7_ = 4.12, *P* = 0.009, Sidak’s *post hoc* test, data not shown). *Grin1*KD prefrontal pyramidal neurons have a significant deficit in dendritic plateau potentials compared to those recorded in brain slices from wild-type littermate mice. This measure confirms a profound physiological impact of insensitivity to glutamate spillover in *Grin1*KD.

### Electrophysiological examination of consequences of adult Grin1 rescue

To identify whether a genetic intervention in adulthood could restore crucial aspects of NMDAR function in *Grin1*KD mice, we tested a tamoxifen-induced Cre-based approach that has previously been shown to increase prefrontal NMDAR radioligand binding and reverse key behavioural deficits ^23^. Briefly, *Grin1*KD mice with loxP sites flanking an insertion Neo cassette were crossed with Cre-ERT2 mice and the adult offspring were treated with tamoxifen (**Fig 4A**). In *Grin1*KD mice with the Cre-ERT2 transgene, tamoxifen induces Cre-mediated excision of the Neo cassette in *Grin1*, restoring full-length mRNA expression and NMDAR levels to ∼60% of wild-type ^23^. These are referred to as *Grin1*rescue mice. In order to ensure equivalent comparison, all 3 genotypes (WT, *Grin1*KD, *Grin1*rescue) were treated with tamoxifen at the same age and for the same time course. Intrinsic electrophysiological properties of prefrontal layer 5 pyramidal neurons including the resting membrane potential, input resistance, capacitance, and action potential amplitude were not significantly different across the tamoxifen-treated, littermate wild-type, *Grin1*KD and *Grin1*rescue mice (**Supplemental Table S2**).

**Figure 4.**
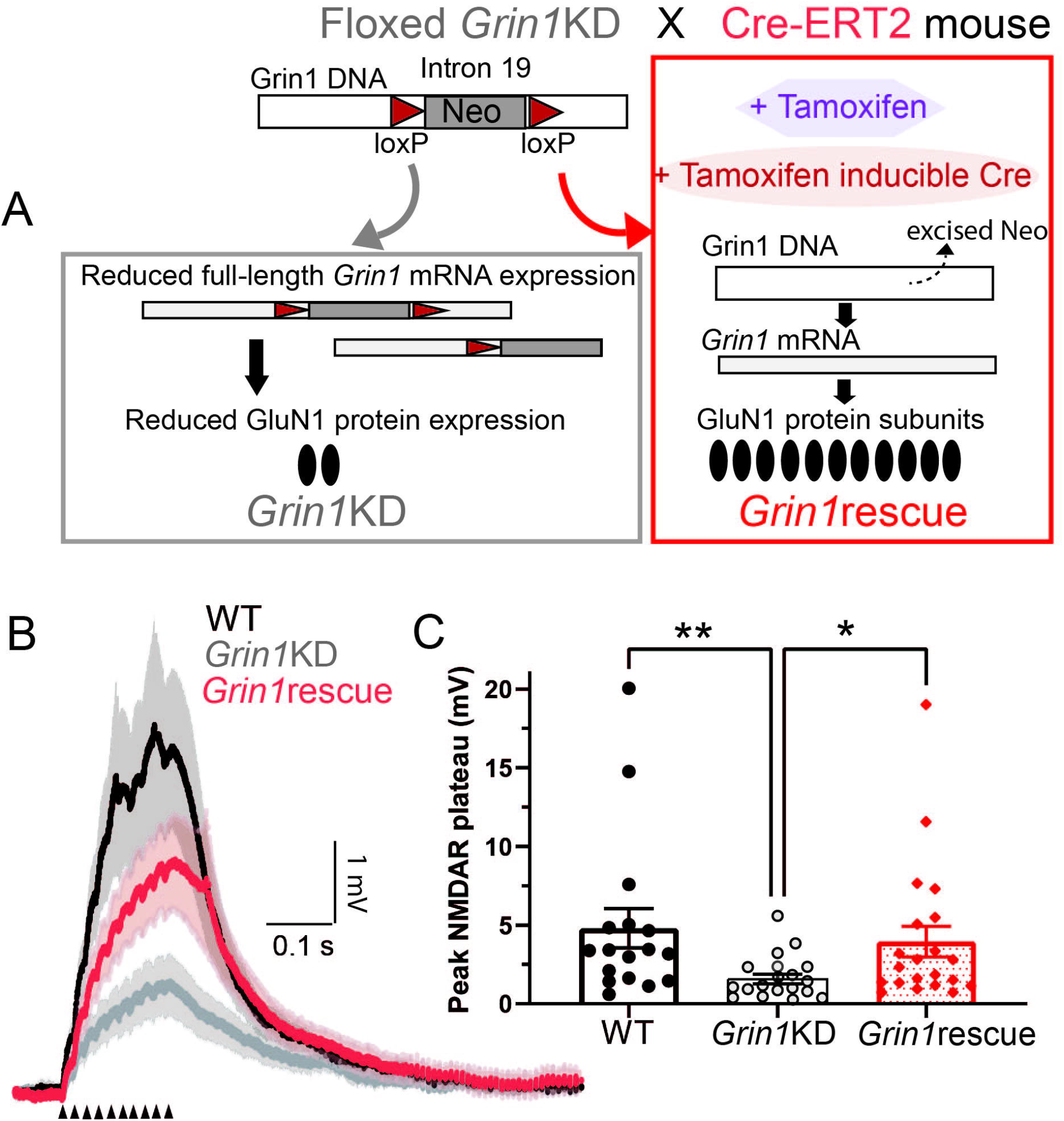
Adult genetic intervention to boost *Grin1* expression restores dendritic plateau potentials. (**A**) *Grin1*rescue schematic illustrates strategy for enhancing *Grin1* expression and increasing NMDAR density in adulthood (adapted from Mielnik and colleagues ^23^). All mice are treated with tamoxifen in adulthood but only in *Grin1*rescue will this treatment trigger Cre expression and lead to the excision of the Neo cassette to increase *Grin1* mRNA, NMDAR radioligand binding, and cognitive performance significantly ^23^. (**B**) Averaged current-clamp recordings of responses to repeated mild stimulation (50 Hz, 10 pulses, 40 μA) in the 3 genotypes of mice all treated with tamoxifen in adulthood: WT (black, *n* = 17), *Grin1*KD (gray, *n* = 18), and *Grin1*rescue (red, *n* = 21). (**C**) Graph illustrates that basal dendrite integration is greatly reduced in *Grin1*KD compared to WT and is restored in the *Grin1*rescue (\*\**P* < 0.01, \**P* < 0.05). Data represented as mean ± SEM.

### Adult intervention rescues dendritic plateau potentials in prefrontal cortex

To identify whether an adult intervention to boost *Grin1* expression can restore dendritic plateau potentials in mice after a lifelong deficit, we examined NMDAR plateau potentials in the three groups of tamoxifen-treated mice. Under these conditions, *Grin1*KD mice again showed significantly smaller NMDAR plateau potentials compared to wild-type mice, but there was a striking increase in the amplitude of the NMDAR plateau potentials in the *Grin1*rescue mice compared to the *Grin1*KD (**Fig 4B**). The distribution of the data prompted nonparametric analysis (Kruskal Wallis test = 11.30, *P* = 0.003; Dunn’s post hoc tests: WT vs *Grin1*KD, *Z* = 3.18, *P* = 0.004; Grin1KD vs *Grin1*rescue, *Z* = 2.55, *P* = 0.032; but no significant difference WT vs *Grin1*rescue, *Z* = 0.76, *P* = 0.99). Correspondingly, dendritic plateau potentials were significantly suppressed by D-APV in both WT and *Grin1*rescue mice (Wilcoxon matched-pairs signed rank test: *P* = 0.016, *n* = 7, data not shown).

Here we show that increasing expression of the obligate NMDAR subunit in adulthood is sufficient to restore dendritic plateau potentials, consistent with the significant behavioural improvement observed previously ^23^. These findings suggest that the boost in *Grin1* expression results in an increase in functional extrasynaptic NMDARs, as illustrated in the working model in **Fig 5.** This work demonstrates the potential for adult treatments to restore NMDAR function critical for signal integration.

**Figure 5.**
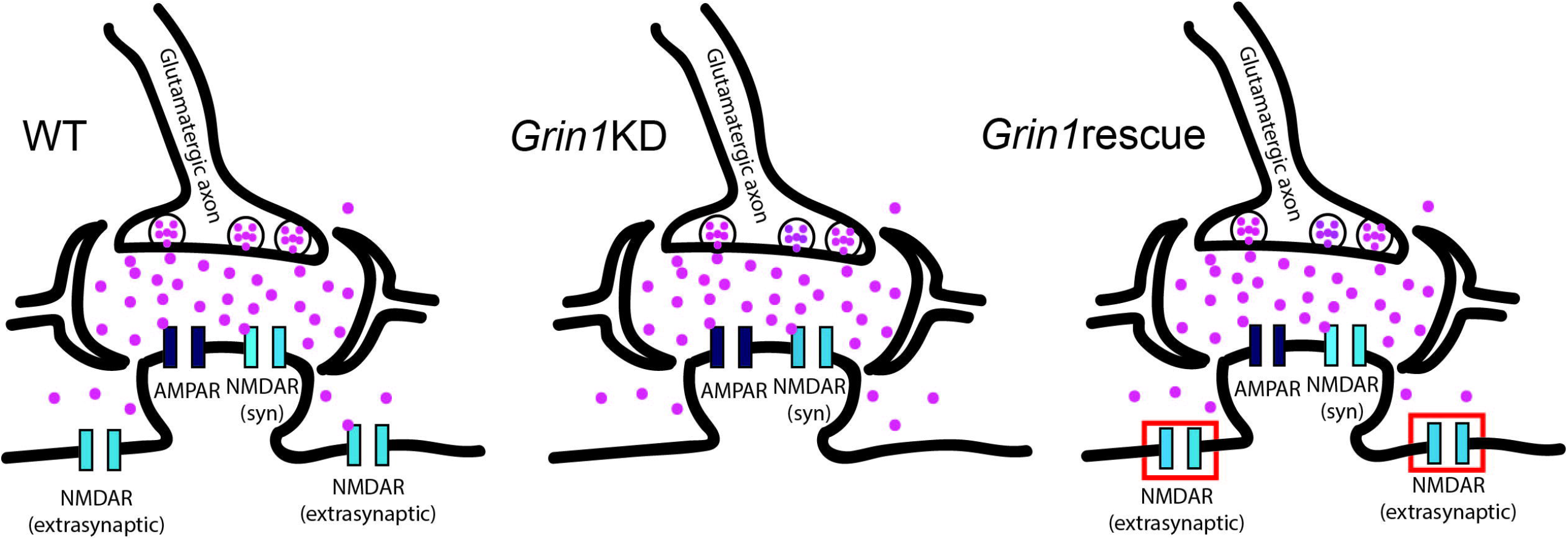
Working model schematics for prefrontal synapses across the three genotypes. In wild-type mice (WT), prefrontal neurons have both synaptic and extrasynaptic NMDARs. In *Grin1*KD mice, there is relative preservation of synaptic NMDARs and disproportionate compromise of extrasynaptic NMDARs. In *Grin1*rescue mice, adult manipulation to boost *Grin1* expression is successful and sufficient to restore extrasynaptic NMDARs needed for dendritic integration of repetitive mild stimuli.

## Discussion

Our data reveal that developmental deficiency in the obligate *Grin1* subunit leads to a profound bias in NMDAR function in the prefrontal cortex. The subpopulation of synaptic NMDARs recruited by mild stimulation shows markedly greater functional preservation than the extrasynaptic receptors recruited by stronger, repetitive, or pharmacological stimuli. To probe the physiological implications of this uneven pattern of NMDAR disruption, we examined dendritic plateau potentials and identified striking deficits in this integrative phenomenon in *Grin1*KD mice. Lastly, we discovered that genetic rescue of *Grin1* expression restores this form of integrative neurophysiology in the mature brain. Our work suggests that, in mice with NMDAR insufficiency, the window for functional improvement remains open into adulthood.

### Broader relevance of this model of NMDAR insufficiency

The *Grin1*KD mouse has been used as a model to study aspects of schizophrenia, autism spectrum disorder, and most recently as a general model for variants in *Grin1* that cause GRIN disorder ^34,35^. *Grin1*KD mice most closely model *Grin1* haploinsufficiency, since they have a genetic modification causing a dramatic reduction in the amount of GluN1 protein and NMDAR without a change in amino acid sequence or in the biophysical properties of the receptor. The *Grin1*KD mouse expresses low levels of the obligate NMDAR subunit with only ∼30% of normal cortical NMDARs, as measured by radioligand binding ^21-23^. Understanding the cellular electrophysiological consequences of this substantial deficit is relevant beyond GRIN disorder, since perturbed NMDAR levels are also a key contributing factor to the symptoms of other neurodevelopmental disorders, including those arising from variants in DLG3, SHANK3, and FMRP ^5,36-40^. Our investigation of *Grin1*KD mice suggest that patients with reduced NMDARs are likely to have a functional deficit in extrasynaptic NMDAR, with a relative preservation of synaptic receptors. Given the historical focus on synaptic NMDAR for neural communication and extrasynaptic receptors for excitotoxicity, it is remarkable that the profound cognitive impairments of *Grin1*KD mice could be attributed to extrasynaptic deficits. It is also striking that rescue experiments in adulthood, which improve executive function and sensory integration ^23^, appear to boost functioning of this extrasynaptic population to restore dendritic plateau potentials, a measure of integrative neurophysiology. This combination of findings urges greater attention to extrasynaptic NMDARs in developmental disorders.

### New perspectives on extrasynaptic NMDARs and their integrative role

Extrasynaptic NMDARs, located perisynaptically ^10^, or non-synaptically on dendritic shafts ^10^, used to be predominantly described in terms of pathology and their role in activating excitotoxic cell death pathways ^41^. However, this view is shifting as growing preclinical research demonstrates the physiological conditions under which extrasynaptic NMDARs are recruited ^15-17^. This recent body of work points to their role in normal brain function via generation of dendritic plateau potentials ^3,4,42^. Extrasynaptic receptors bind the small amount of glutamate that escapes the synapse, to become ‘primed’ and ready for rapid activation by subsequent depolarizing input(s). NMDARs on small dendritic branches are thus positioned to detect the activation of multiple synapses close together in space and time. Such temporal and spatial integration is required to generate dendritic plateau potentials ^2,7,18,19^. These NMDAR-mediated integration events trigger burst firing ^7,19^, a robust neuronal response ^32,33^, thought to be essential for behaviour-evoked network activity ^4,20,43^. Our results indicate that developmental disorders with reduced NMDARs are likely to have compromised neurophysiological integration resulting from disrupted extrasynaptic NMDAR population. Intriguingly, an adult intervention yielding an increase in *Grin1* expression and NMDAR radioligand binding ^23^ (to ∼60% of wild-type), restores the neurophysiological phenomenon of dendritic plateau potentials. This integrative recovery is consistent with the marked improvement of cognitive performance observed after treatment in adulthood ^23^.

### Subcompartment-specific NMDAR alterations: potential mechanisms and caveats

Disparate functional consequences across NMDAR populations have been observed in response to different perturbations ^44-48^. Research in cell systems demonstrates that NMDARs move between synaptic and extrasynaptic compartments upon pharmacological manipulation ^44-47^, or exposure to antibodies from people with anti-NMDAR encephalitis ^49^. Receptor trafficking, however, is not the only path to achieve divergent functional outcomes for synaptic and extrasynaptic NMDAR populations. Multiple mechanisms for functional NMDAR enhancement display compartmental specificity, including post-translational modification pathways ^50,51^, co-agonism ^52-54^, and mechanisms of receptor desensitization ^55-57^. The functional preservation of synaptic NMDAR responses in *Grin1*KD mice may therefore be caused by multiple complex mechanisms, and not necessarily reflect wild-type levels of receptor density in this compartment ^23^.

While NMDARs are the focus of a large body of work in models of neurodevelopmental disorders, many characterizations use relatively strong stimuli under conditions where ‘synaptic’ measures may inadvertently include a broader population. Here, we pursued carefully calibrated electrical stimulation under several conditions to isolate synaptic NMDARs from their extrasynaptic counterparts. Our strategy was adopted due to the inherent challenges in separating these contributions with pharmacological tools ^58,59^. This problem is particularly difficult to overcome in the prefrontal cortex, where synaptic and extrasynaptic NMDARs show a high degree of overlap in molecular composition and pharmacological affinities ^60,61^, complicating specific manipulations. Differentiating synaptic and extrasynaptic NMDAR populations remains a challenging, but increasingly important, focus for future work into the mechanisms of cognitive compromise arising from NMDAR insufficiency.

### Clinical relevance and future implications

Current treatments for cognitive disability arising from genetic disruption of NMDARs focus on supportive therapies because it is assumed that lifting cognitive restrictions hard-wired by abnormal brain development is impossible. However, this assumption has recently been challenged. Promising preclinical data ^23,62,63^ suggest the potential for cognitive improvement, even when intervention is delayed until adulthood. If adult treatments are to be seriously pursued, it is essential to appreciate what neural components are functionally compromised and what may be preserved. Here, we address a critical knowledge gap about the specific cellular and circuit mechanisms by which genetic NMDAR disruption impairs cognitive function. We demonstrate that two important NMDAR subpopulations do not suffer equal consequences from genetic disruption of the obligate subunit *Grin1*. Extrasynaptic NMDARs are disproportionately compromised with resulting disruption of the integrative capacity required for the generation of dendritic plateau potentials. This deficit, strikingly, proves amenable to rescue by intervention in adulthood. Developing effective treatments for the cognitive impairments caused by NMDAR disruption requires the identification of the most efficient targets. Our discovery underscores the need for research into additional approaches to safely enhance extrasynaptic NMDAR functioning. Overall, our findings suggest that deficient integrative mechanisms are amenable to improvement, even with adult intervention.

## Acknowledgements

This work was generously supported by the Canadian Institutes of Health Research (EKL, PJT-178372, EKL MOP-89825; AJR MOP-119298), Simons Foundation for Autism Research (AJR), CureGRIN Research Foundation (AJR), the Canada Research Chairs program (EKL, Canada Research Chair in Developmental Cortical Physiology), a Pathway Award from the University of Toronto Faculty of Medicine (EKL), as well as multiple fellowships from the Ontario Graduate Scholarship programs (SV, MAB, CAM). We gratefully thank Wendy Horsfall, Duong Chu, Janice McNabb, and Marija Milenkovic for expert technical assistance.

