## Supplemental Table S1 and Table S2, Suppl Methods for "Deficits in integrative NMDA receptors caused by *Grin1* disruption can be rescued in adulthood"

Your manuscript submission has been assigned the following tracking number: **NPP-22-1263**. Please quote this number in any correspondence with the editorial office.

**Author Instructions**

- The manuscript submission process consists of 4 primary tasks: 1. Files, 2. Manuscript Information, 3.Validate, 4. Submit. You will need to complete the primary tasks in the correct order.
- You will have the opportunity to make changes to your submission until you click the 'Approve Manuscript' button on the 'Approve Manuscript' tab.
- To save a draft version of your manuscript to complete at a later stage, click on the 'Save and Exit' button. You will then return to your author desktop
- **Still confused?** Click here for further instructions on how to complete the manuscript submission process.
- **Click** here for a checklist containing items to confirm while you submit.
- NOTE: * indicates a required Field

1. Files | 2. Manuscript Information | **3. Validate** | 4. Submit

a) Approve Files    b) Approve Manuscript Details

*No errors found.*

| | |
|---|---|
| **Title** * | Deficits in integrative NMDA receptors caused by Grin1 disruption can be rescued in adulthood<br>Change |
| **Abstract** * | Glutamatergic NMDA receptors (NMDAR) are critical for cognitive function, and their reduced expression leads to intellectual disability. Since subpopulations of NMDARs exist in distinct subcellular environments, their functioning may be unevenly vulnerable to genetic disruption. Here, we investigate synaptic and extrasynaptic NMDARs on the major output neurons of the prefrontal cortex in mice deficient for the obligate NMDAR subunit encoded by Grin1 and wild-type littermates. With whole-cell recording in brain slices, we find that single, low-intensity stimuli elicit surprisingly-similar glutamatergic synaptic currents in both genotypes. By contrast, clear genotype differences emerge with manipulations that recruit extrasynaptic NMDARs, including stronger, repetitive, or pharmacological stimulation. These results reveal a disproportionate functional deficit of extrasynaptic NMDARs compared to their synaptic counterparts. To probe the repercussions of this deficit, we examine an NMDAR-dependent phenomenon considered a building block of cognitive integration, basal dendrite plateau potentials. Since we find this phenomenon is readily evoked in wild-type but not in Grin1-deficient mice, we ask whether plateau potentials can be restored by an adult intervention to increase Grin1 expression. This genetic manipulation, previously shown to restore cognitive performance in adulthood, successfully rescues electrically-evoked basal dendrite plateau potentials after a lifetime of NMDAR compromise. Taken together, our work demonstrates NMDAR subpopulations are not uniformly vulnerable to the genetic disruption of their obligate subunit. Furthermore, the window for functional rescue of the more-sensitive integrative NMDARs remains open into adulthood.<br>Change |

| | |
|---|---|
| **b) Authors *** | Sridevi Venkatesan (University of Toronto) (Contrib #1)<br><br>Mary A Binko (University of Toronto) (Contrib #2)<br><br>Catharine A Mielnik (University of Toronto) (Contrib #3)<br><br>Amy J Ramsey (University of Toronto) (Contrib #4)<br><br>Evelyn K Lambe (University of Toronto) (Corr)<br><br>Change |
| **Techniques *** | Patch clamp<br>Transgenic mice<br>Change |
| **Category *** | Electrophysiology<br>Change |
| **Subject Terms *** | Subject Terms: Biological sciences/Neuroscience/Ion channels in the nervous system, Biological sciences/Physiology/Neurophysiology, Health sciences/Medical research/Preclinical research,<br>Change |
| **Social Media** | *If you would like to supply our social media editor with a "tweet", which may or may not be used or edited at the discretion of the journal, please insert the message here (280 characters):*<br>Which NMDA recptors?! Grin1 deficiency leads to selective loss of extrasynaptic #NMDAR, compromising dendritic plateau potentials. Adult intervention can bring them back! #CureGRIN #autism<br><br>*Please enter the Twitter handle of you and/or your co-authors.*<br>@LambeLab @sridevi292 @camielnik @ajr937<br>Change |
| **Collection of papers *** | *On our website, NPP features a collection of papers entitled "Highlighting Research on Health Disparities". If this manuscript is accepted, should the editors consider it for inclusion in this collection?*<br>No<br><br>*If you answered yes, please provide a 1-sentence justification:*<br>[Empty]<br>Change |
| **Clinical Trials** | *NPP follows the International Committee of Medical Journal Editors definition of clinical trials.*<br><br>*According to the definition, a clinical trial is a research study in which one or more human subjects are prospectively assigned to one or more interventions (which may include placebo or other control) to evaluate the effects of those interventions on health-related biomedical or behavioral outcomes. Note: Basic experimental studies involving humans subjects can be considered a clinical trial if subjects are prospectively assigned to an intervention and the study evaluates the effects of the intervention. More info here.*<br><br>*If your manuscript does not involve any human participants, answer NO to the following five questions*<br><br>*1. Does your study involve human participants?*<br>No<br><br>*2. Are the human participants prospectively assigned to an intervention?*<br>No<br><br>*3. Is the study designed to evaluate the effect of the intervention on the human?*<br>No<br><br>*4. Is the effect being evaluated a health related biomedical (brain or peripheral measure) or behavioral outcome?*<br>No<br><br>*If you have answered yes to all 4 of these questions, your study meets the definition of a clinical trial. Therefore, you are required to include the following three mandatory items with your submission materials:* |

| | |
|---|---|
| | *1. A completed CONSORT checklist.*<br>*2. The clinical trial registry number in the manuscript.*<br>*3. The CONSORT Flow Diagram.*<br>*Failure to include all three mandatory items with your submission will result in a delay of the processing of your manuscript. If any concerns are raised by editorial staff about manuscript compliance, your submission will be sent to the senior editors for review.*<br>*5. I confirm that the study involves human participants and I have included in my submission materials all three of the following: 1) a completed CONSORT Checklist; 2) the registry number in the manuscript; 3) the CONSORT Flow Diagram*<br>No<br>Change |
| **Research Square author dashboard *** | I understand that my manuscript and associated personal data will be shared with Research Square for the delivery of the author dashboard.<br>Change |
| **Conflict of Interest Statement *** | There is **NO** conflict of interest to disclose.<br>Change |
| **Clinical Trial *** | Clinical Trial: No<br>Change |
| **Archiving Mandates *** | Springer Nature permits the authors of all original research papers published in Nature Portfolio titles or academic journals on nature.com to self-archive the author's accepted manuscript (AAM) in an institutional or funder repository, where it can be made publicly accessible 6 months after publication in accordance with our self-archiving policy.<br>Several funders require deposition of manuscripts to PubMed Central or Europe PubMed Central. To enable deposition and compliance with these requirements, Springer Nature offers a free manuscript deposition service for original research papers supported by a number of PMC/EPMC participating funders. With your authorization, we can deposit manuscripts in PMC/Europe PMC on your behalf. We encourage you to take advantage of this service if you or any of your co-authors are funded by one of the PMC/Europe PMC participating funders listed on our website.<br>Please select from the following options:<br>I do not wish Springer Nature to archive this manuscript if accepted, or this service is not applicable to me and my co-authors. If my or my co-authorsâ€™ funder mandates self-archiving, I understand that it is my responsibility to archive in accordance with its policies.<br>Change |
| **Authorship *** | Yes<br>Change |
| **Manuscript Comment** | *Additional Comments:*<br>[Empty]<br>Change |
| **Previous Interactions** | Change |
| **Color Art *** | Yes, my manuscript contains figures that must be published in color in the PDF/print hard copy versions, and I understand, if my article is accepted, I will be invoiced to pay the associated color charges.<br>Comments:<br>Fig 2A, Fig 3A, Fig 4, Fig 5<br>Change |
| **VAT Number** | [Empty]<br>Change |

| | |
|---|---|
| **Purchase Order** | [Empty]<br>Change |
| **Suggested Reviewers to Include** | *Suggested Reviewers to Include*<br>Lynn Raymond,, UBC<br>Wen Jun Gao, , Drexel<br>Suzanne Zukin, , Einstein<br>Tak Pan Wong, , McGill<br>Brady Maher, , Johns Hopkins<br>Helen Barbas, , Boston University<br>Change |
| **Suggested Reviewers to Exclude** | *Suggested Reviewers to Exclude*<br>Steven Siegel,, USC<br>Jon Johnson,, UPitt<br>Change |

Back | Save and Exit | Save and Continue

Not yet completed
Required fields are incomplete
Complete

eJournalPress

**tracking system home** | **author instructions** | **reviewer instructions** | **help** | **tips** | **logout** | **journal home** | **terms of use**
**privacy policy** | **cookie policy** | **manage cookies**
